# Development of potent humanized TNFα inhibitory nanobodies for therapeutic applications in TNFα-mediated diseases

**DOI:** 10.1101/2025.02.07.637018

**Authors:** Tao Yin, Aubin Ramon, Matthew Greenig, Pietro Sormanni, Luciano D’Adamio

## Abstract

Tumor necrosis factor-alpha (TNFα) is a key pro-inflammatory cytokine implicated in the pathogenesis of numerous inflammatory and autoimmune diseases, including rheumatoid arthritis, inflammatory bowel disease, and neurodegenerative disorders such as Alzheimer’s Disease. Effective inhibition of TNFα is essential for mitigating disease progression and improving patient outcomes. In this study, we present the development and comprehensive characterization of potent humanized TNFα inhibitory nanobodies (TNFI-Nbs) derived from camelid single-domain antibodies. In silico analysis of the original camelid nanobodies revealed low immunogenicity, which was further reduced through machine-learning-guided humanization and developability optimization. The two humanized TNFI-Nb variants we developed demonstrated exceptional anti-TNFα activity, achieving IC_50_ values in the picomolar range. Binding assays confirmed their high affinity for TNFα, underscoring robust neutralization capabilities. These TNFI-Nbs present valid alternatives to conventional monoclonal antibodies currently used in human therapy, offering potential advantages in potency, specificity, and reduced immunogenicity. Our findings establish a solid foundation for further preclinical development and clinical translation of TNFα-targeted nanobody therapies in TNFα-mediated diseases.

## Introduction

TNFα is a key pro-inflammatory cytokine implicated in the pathogenesis of numerous inflammatory and autoimmune diseases, including rheumatoid arthritis, inflammatory bowel disease, and neurodegenerative disorders such as Alzheimer’s Disease. Effective inhibition of TNFα is essential for mitigating disease progression and improving patient outcomes. Biologic TNFα inhibitors (TNFI), including conventional monoclonal antibodies, have revolutionized the treatment of peripheral TNFα-mediated conditions (1-5). However, conventional antibodies face limitations such as large molecular size, potential immunogenicity, and restricted tissue penetration.

Nanobodies (Nbs), or single-domain antibodies derived from camelid species, offer a promising alternative to conventional antibodies (6, 7). These molecules consist of a single monomeric variable antibody domain, featuring smaller paratopes that can access functional pockets of target proteins more efficiently. This results in enhanced binding specificity, affinity, and inhibitory efficiency, allowing nanobodies to recognize and bind epitopes often inaccessible to traditional antibodies, thereby broadening therapeutic targets. Additionally, their smaller size (∼12 kDa compared to ∼150 kDa for conventional antibodies) reduces the potential for immunogenicity and facilitates better tissue penetration. Nanobodies also exhibit remarkable stability under various physiological conditions, including extreme pH and temperature, which is advantageous for both storage and administration. However, further improvements in stability are necessary to maximize their utility in peripheral inflammatory therapies, ensuring sustained therapeutic action and minimizing the need for frequent dosing.

Beyond peripheral inflammatory conditions, nanobodies hold significant potential for the treatment of central nervous system (CNS) diseases. Conventional antibodies have limited blood-brain barrier (BBB) permeability, restricting their effectiveness in targeting CNS pathologies. In contrast, the smaller size and robust structural stability of nanobodies may facilitate better BBB penetration (8, 9), enhancing their therapeutic reach within the brain. Moreover, their inherent ability to be engineered for improved BBB permeability opens avenues for innovative delivery strategies, such as receptor-mediated transcytosis via transferrin receptor 1 (TfR1) (10-14), potentially increasing their efficacy in treating neurodegenerative disorders.

In this study, we present the development and comprehensive characterization of potent humanized TNFα inhibitory nanobodies (TNFI-Nbs) derived from camelid single-domain antibodies. By leveraging advances in silico analysis and machine-learning-guided mutagenesis, we optimized the framework of TNFI-Nbs to maximize humanness and solubility while retaining and slightly improving affinity and for TNFα. Our findings demonstrate that these engineered nanobodies exhibit exceptional anti-TNFα activity and possess the structural and functional attributes necessary for potential therapeutic applications in both peripheral inflammatory and CNS diseases. Furthermore, we explore strategies to enhance BBB permeability through targeted transcytosis mechanisms, positioning TNFI-Nbs as versatile and effective tools against TNFα-mediated pathologies.

## RESULTS

### Generation of anti-human TNFα nanobodies

To generate nanobodies (Nbs) specifically targeting human TNFα, one llama and one alpaca were immunized with active trimeric human TNFα (Acro Biosystems, TNA-H5228) to elicit a robust immune response. Serum titers from the immunized animals were monitored and quantified using ELISA on antigen-coated plates to confirm the production of high-affinity antibodies. Peripheral blood mononuclear cells (PBMCs) were isolated and cDNAs encoding the VHH domains of Nbs were cloned into the pADL-20c phagemid vector and transformed into *E. coli* TG1 cells for phage display.

To enrich nanobody clones with strong binding to human TNFα, multiple rounds of panning were performed on antigen-coated plates, a process designed to selectively capture high-affinity clones while removing non-specific or weak binders. After enrichment, 470 individual nanobody clones were expressed in *E. coli*, and His-tagged nanobodies were recovered from the periplasmic fraction via osmotic shock. These nanobodies were screened for their ability to bind human TNFα by ELISA, leading to the identification of α-TNFα nanobodies (α-TNFα-Nbs) with strong antigen interaction. Sequencing of ELISA-positive clones revealed 100 unique α-TNFα-Nb sequences, each representing a distinct nanobody (**Table 1**). These 100 unique nanobodies were then expressed in bacterial cultures and purified using affinity chromatography. The purified α-TNFα-Nbs underwent further functional characterization.

**Table 1.**
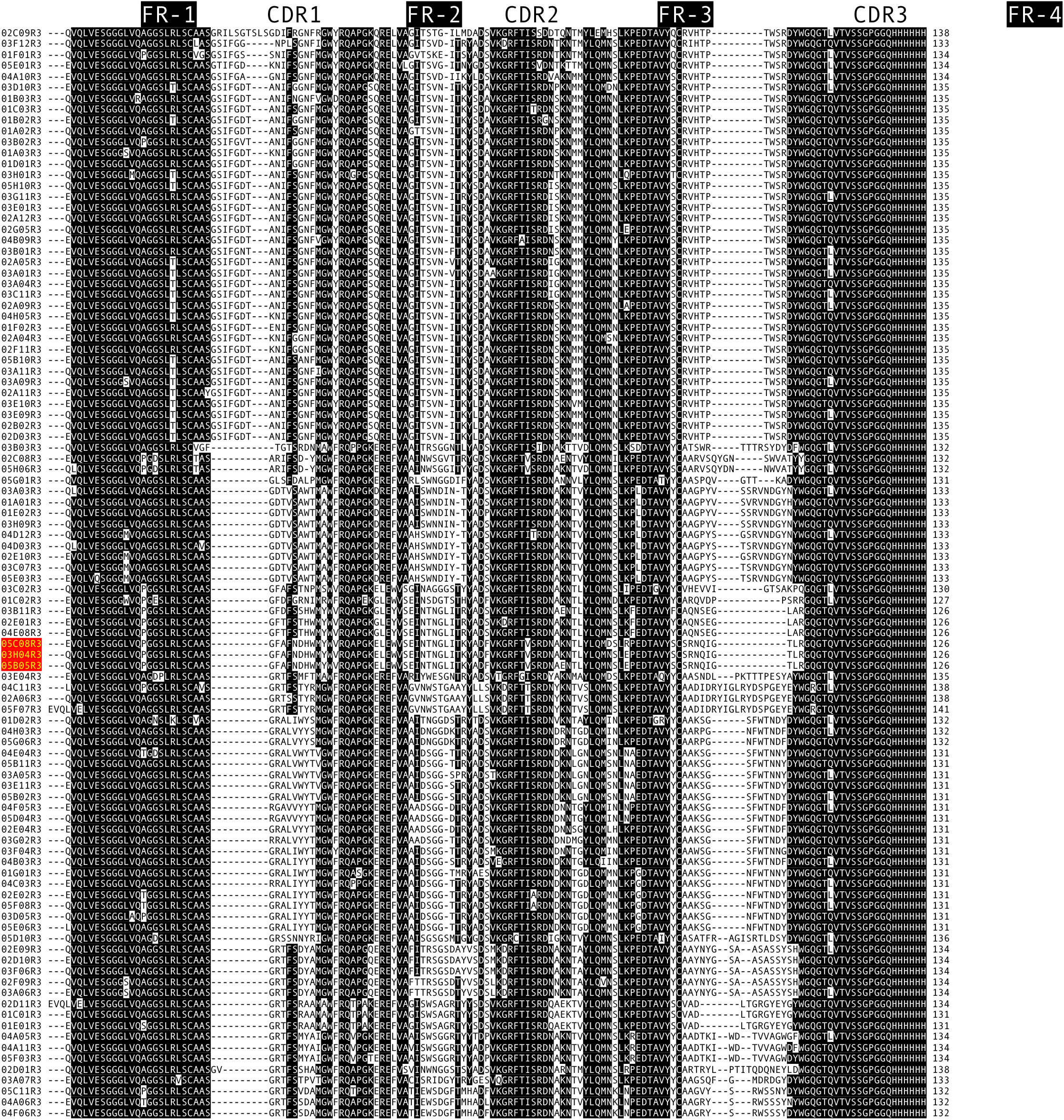
Amino Acid Sequences of 100 Unique α-TNFα-Nbs Isolated from an Alpaca and a Llama Immunized with Active Trimeric Recombinant Human TNFα. The α-TNFα-Nbs are listed in order of complementarity-determining region (CDR) similarity and sequence identity. The three α-TNFα-Nbs highlighted in red represent the top candidates, demonstrating the highest TNFα inhibitory activity in subsequent functional assays. Framework regions (FR) are highlighted in black.

The above experiments were outsourced to Abcore, a contract research organization (CRO) specializing in the generation of custom antibodies, including nanobodies, for research and therapeutic applications.

### Identification of α-TNFα nanobodies with high inhibitory activity against human TNFα (TNFINbs)

The 100 purified α-TNFα-Nbs were assessed for their ability to inhibit cytotoxicity induced by active trimeric human TNFα. Initial screening utilized WEHI-13VAR cells, a mouse line that serves as a sensitive bioassay for both rodent and human TNFα. WEHI-13VAR cells were incubated for 24 hours with 0.25 ng/mL active trimeric human TNFα and 1 μg/mL Actinomycin-D, either alone or in the presence of 100 nM of each α-TNFα-Nb. Cytotoxicity was measured using the Cell Counting Kit-8, comparing cell viability to that of cells treated with Actinomycin-D alone. Eleven α-TNFα-Nbs that achieved at least 60% inhibition of human TNFα–induced cell death at 100 nM concentration (data not shown) were selected for further evaluation. In the secondary screening, these selected α-TNFα-Nbs underwent 2-fold serial dilutions ranging from 100 nM to 0.76294 pM. Secondary screening identified three nanobodies (05C08R3, 05B05R3, and 03H04R3) with strong TNFα inhibitory activity, exhibiting IC_50_ values of 95.78 pM (95% confidence interval [CI]: 78.24–116.4 pM), 219.3 pM (95% CI: 178.7–268.3 pM), and 175 pM (95% CI: 136–222.7 pM), respectively (**Figure 1A**). Consequently, these α-TNFα nanobodies were renamed TNFI-Nb1, TNFINb2, and TNFI-Nb3.

**Figure 1.**
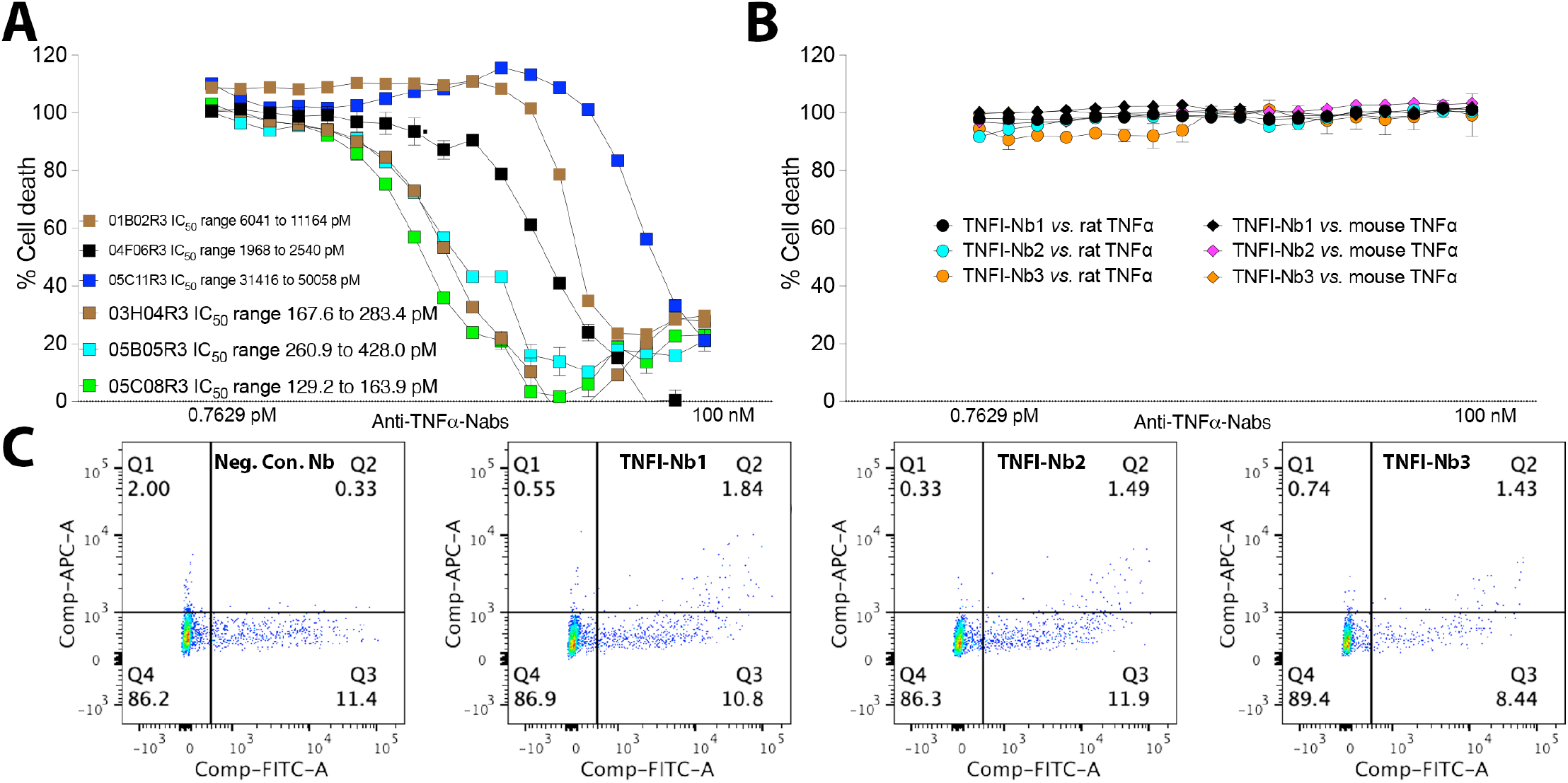
TNFI-Nb1, TNFI-Nb2, and TNFI-Nb3 exhibit potent, human-specific TNFα inhibition and bind membrane-bound TNFα. **A**) The inhibitory activity of the selected α-TNFα-Nbs was assessed in a secondary screening using 2-fold serial dilutions ranging from 100 nM to 0.76294 pM. Results are presented as mean ± SEM of triplicate measurements and analyzed using the “Inhibitor vs. Normalized Response” model in GraphPad Prism 10 software. IC50 values, representing the concentration required to achieve 50% inhibition, are expressed in picomolar (pM) units. The three α-TNFα-Nbs with the highest inhibitory activity were named TNFI-Nb1 (05C08R3), TNFI-Nb2 (05B05R3), and TNFI-Nb3 (03H04R3). **B**) TNFI-Nb1, TNFI-Nb2, and TNFI-Nb3 do not inhibit rat and mouse TNFα. The inhibitory activities of TNFI-Nb1, TNFI-Nb2, and TNFI-Nb3 against active trimeric rat and mouse TNFα were evaluated under the same assay conditions used for human TNFα (see Figure 1). The results demonstrate that none of the three TNFI-Nbs inhibited rat or mouse TNFα at the concentrations tested (2-fold serial dilutions ranging from 100 nM to 0.76 pM). Data are presented as mean ± SEM from triplicate measurements and were analyzed using the “Inhibitor vs. Normalized Response” model in GraphPad Prism 10 software. **C**) HEK293 cells were transfected with a vector expressing human TNFα plus EGFP. transfected cells were incubated with 40nM α-TNFα-Nb, followed by an anti-His-APC antibody (R&D Systems, IC050A). Staining with a nanobody that does not bind human TNFα, along with anti-His-APC, was used as a negative control.

Sequence analysis of the three selected nanobodies revealed identical complementarity-determining regions (CDRs), with sequence differences confined to framework regions (FR) 1 and 3 (**Table 2**). Consequently, these three TNFI-Nbs belong to the same family, and their identical CDRs account for their similar high TNFI activity. This conclusion is further supported by the observation that these three nanobodies were the only ones among the 100 tested that belonged to this family (**Table 1**).

**Table 2.**
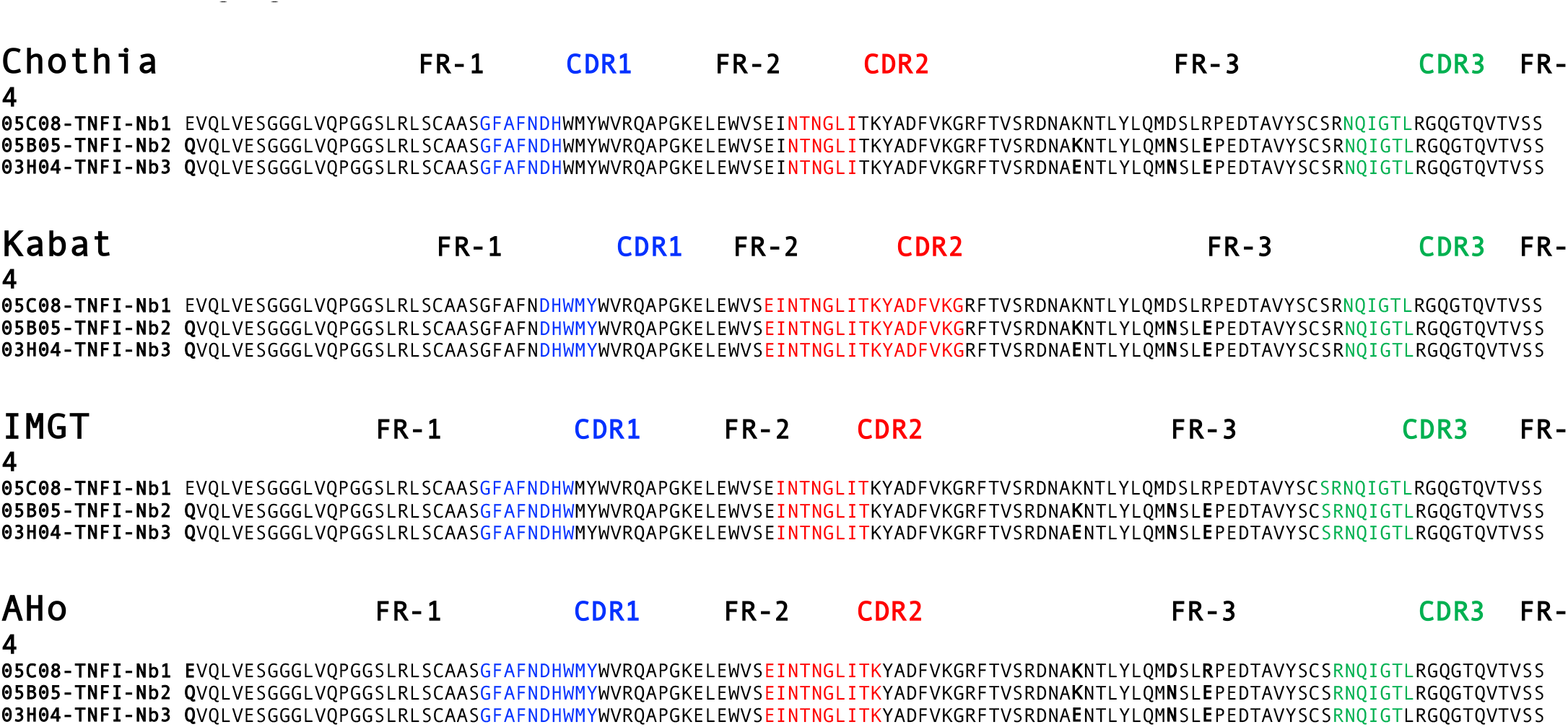
Alignment of Amino Acid Sequences of TNFI-Nb1 (05C08R3), TNFI-Nb2 (05B05R3), and TNFI-Nb3 (03H04R3). The amino acid sequences of TNFI-Nb1, TNFI-Nb2, and TNFI-Nb3 are aligned to illustrate their structural similarity. All three nanobodies share identical CDR1, CDR2, and CDR3 sequences across all numbering systems. Sequence differences, highlighted in bold, are confined to the FR-1 and FR-3 regions. The alignment accommodates multiple numbering schemes, including Chothia (structural loop-based), Kabat (sequence variability-based), IMGT (universal immunogenetics standard), and AHo (comprehensive immunoglobulin-like domain framework). This comprehensive approach ensures consistency and facilitates comparisons of sequence and structure across diverse analytical contexts.

To further characterize the specificity of TNFI-Nb1, TNFI-Nb2, and TNFI-Nb3, we evaluated their ability to inhibit cytotoxicity induced by rat and mouse TNFα. Utilizing the same assay conditions described for human TNFα inhibition, WEHI-13VAR cells were incubated for 24 hours with active trimeric rat TNFα or mouse TNFα in the presence of each nanobody. The nanobodies were tested at a concentration of 100 nM, followed by 2-fold serial dilutions ranging from 100 nM to 0.76294 pM (**Figure 1B**). Our results demonstrate that none of the three TNFI-Nbs were able to inhibit cytotoxicity induced by either rat or mouse TNFα at any of the concentrations tested. This lack of inhibitory activity underscores the high specificity of these nanobodies for human TNFα. This selectivity for human TNFα makes standard rodent models unsuitable for evaluating the biological activities of TNFI-Nb1, TNFI-Nb2, and TNFI-Nb3. Therefore, humanized animal models or in vitro systems expressing human TNFα are necessary to investigate their therapeutic potential and mechanisms of action. These models will enable accurate assessment of the nanobodies’ efficacy and safety, providing essential data for advancing to clinical trials.

Finally, we evaluated whether TNFI-Nb1, TNFI-Nb2, and TNFI-Nb3 can bind to membrane-bound TNFα. To achieve this, HEK293 cells were transfected with a construct expressing human TNFα and EGFP to label the transfected cells. As shown in **Figure 1C**, flow cytometry (FACS) analysis revealed that all three TNFI-Nbs bound to the transfected cells, demonstrating their ability to recognize and interact with membrane-bound TNFα.

### Production of TNFI-Nb1 in CHO-S cells

Chinese Hamster Ovary (CHO) cells are a preferred system to produce therapeutic proteins due to their ability to generate complex proteins with proper folding, posttranslational modifications, and biological activity similar to native human proteins. Another significant advantage of using CHO cells is their ability to produce proteins with significantly lower endotoxin levels compared to bacterial expression systems, enhancing the safety and suitability of the final product for therapeutic use. Therefore, this system minimizes the risk of immunogenicity by ensuring the produced proteins are more likely to be recognized as “self” by mammalian immune systems. Given their long history of regulatory approval for numerous therapeutic proteins, CHO cells provide a reliable platform for biopharmaceutical manufacturing.

To determine the optimal production system for TNFI-Nb1, we compared protein production efficiency and quality between *E. coli* and CHO-S cells. CHO-S cells produced 16.52 mg of TNFI-Nb1 from a 100 mL culture at a concentration of 1.18 mg/mL, with a purity of ≥95% as assessed by SDS-PAGE and 96% by SEC-HPLC. Endotoxin levels in the CHO-S-derived protein were ≤0.1 EU/mg, well below acceptable thresholds for therapeutic use. It is important to note that these substantial, yet not exceptional, production levels were achieved through transient transfection without selecting high-producing stable clones or implementing further process optimization. In contrast, *E. coli* yielded only 2.1 mg of TNFI-Nb1 from a 1 L culture at a concentration of 0.35 mg/mL, with lower purity (≥90% by SDS-PAGE) and significantly higher endotoxin levels (≤10.9 EU/mg). These results clearly demonstrate the superiority of CHO-S cells in terms of yield, purity, and endotoxin levels.

Based on these findings, we produced TNFI-Nb2 and TNFI-Nb3 in 100 mL CHO-S cell cultures. TNFI-Nb2 was obtained at a yield of 14.26 mg (0.92 mg/mL) with a purity of ≥90% as assessed by SDS-PAGE and 94% by SEC-HPLC, and endotoxin levels ≤0.1 EU/mg. Similarly, TNFI-Nb3 was produced at 13.64 mg (0.88 mg/mL) with a purity of ≥95% by SDS-PAGE and 95% by SEC-HPLC, and endotoxin levels ≤0.1 EU/mg. Protein production was outsourced to GenScript.

### Surface Plasmon Resonance (SPR) analysis of TNFI-Nb1-3 binding to human TNFα

To determine the binding affinity of TNFI-Nb1, TNFI-Nb2, and TNFI-Nb3 to human TNFα, we conducted SPR analysis. The purity of all three nanobodies—TNFI-Nb1 (05C08R3,VHH1), TNFI-Nb2 (05B05R3, VHH2), and TNFI-Nb3 (03H04R3, VHH3)—as well as human TNFα, was confirmed to exceed 90%, ensuring highquality reagents for subsequent assays. Kinetic analyses were performed using Cytiva’s Biacore 1K SPR system and Nicoya’s OpenSPR-XT instrument with Carboxyl CM5 sensors. Assay optimization encompassed the screening of various immobilization conditions, buffer formulations, and regeneration protocols to establish a reliable and reproducible measurement framework. Optimal immobilization was achieved using the Nbs the ligands at a concentration of 2.5 μg/mL in 10 mM acetate buffer (pH 4.5), resulting in appropriate response maxima (Rmax) for each nanobody and ensuring sufficient signal for accurate kinetic fitting. Buffer optimization identified 1x PBS supplemented with 3 mM EDTA and 0.05% v/v Surfactant P20 as the optimal running buffer, effectively minimizing non-specific binding and maximizing signal-to-noise ratios.

Multi-cycle kinetic analysis revealed that all three nanobodies interacted with TNFα following a 1:1 binding model, demonstrating high affinity in the picomolar range (**Figure 2A**). TNFI-Nb2 exhibited the highest affinity with an equilibrium dissociation constant (KD) of 77 pM, primarily driven by its slower off-rate (kd (s^-1^) = 2.87E-05 ±4.6E-06). TNFI-Nb1 displayed a KD of 219 pM, while TNFI-Nb3 showed a KD of 392 pM (**Figure 2B**). Iso-affinity analysis further confirmed that TNFI-Nb2’s superior affinity was mainly due to its slower dissociation rate, distinguishing it from TNFI-Nb1 and TNFI-Nb3 (**Figure 2C**). These findings collectively highlight the potent TNFα inhibitory capabilities of the nanobody panel, underscoring their potential as effective therapeutic agents. These SPR results confirm the potent and specific binding of TNFI-Nb1, TNFI-Nb2, and TNFI-Nb3 to human TNFα, validating their potential as effective therapeutic agents. SPR experiments were outsourced to Rapid Novor.

**Figure 2.**
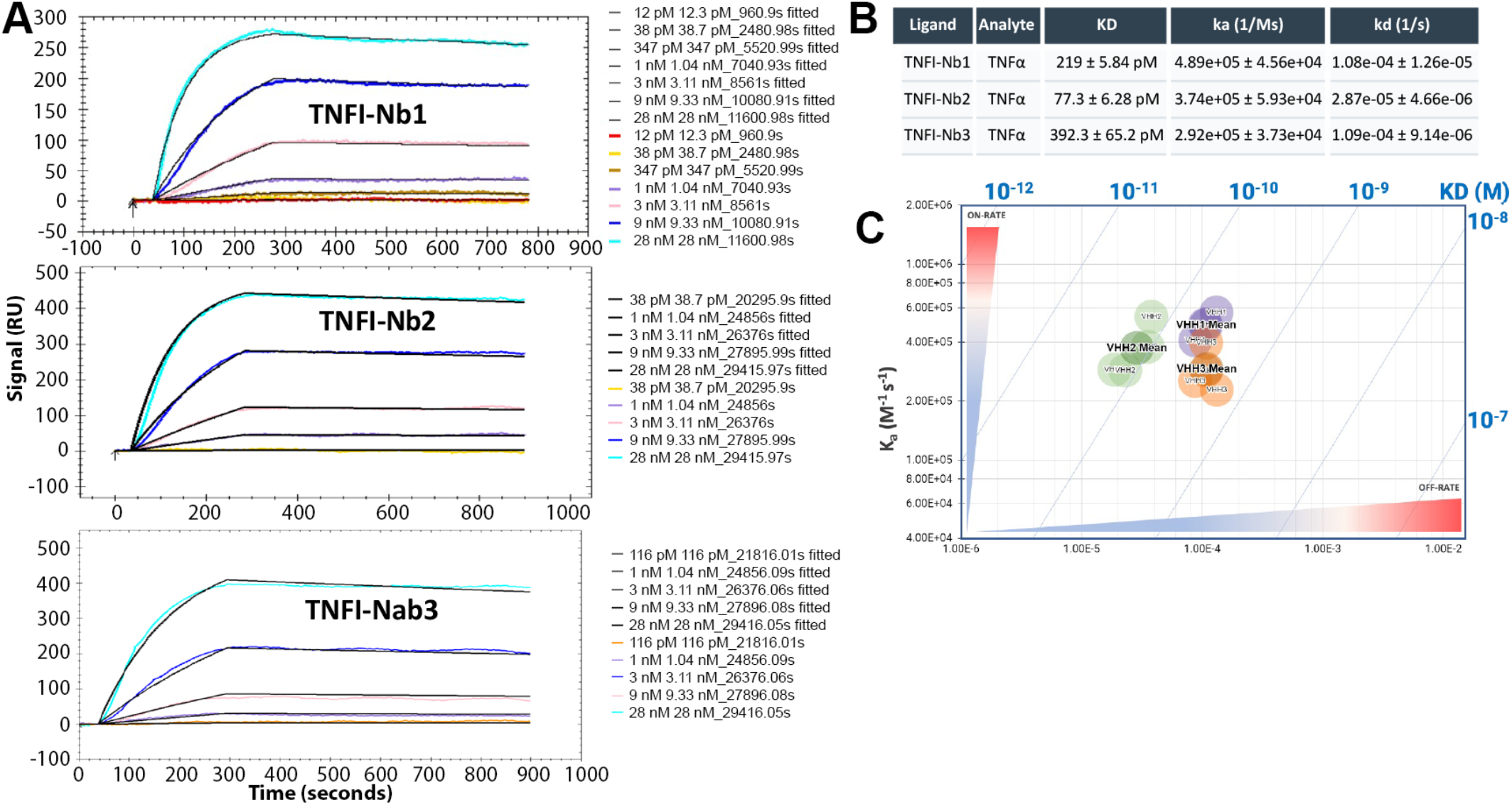
Kinetics of TNFα binding to immobilized TNFI-Nb1, TNFI-Nb2 and TNFI-Nb3. Kinetic analysis of TNFα binding to nanobodies TNFI-Nb1 (VHH1), TNFI-Nb2 (VHH2), and TNFI-Nb3 (VHH3) directly immobilized on Carboxyl sensors via amine coupling. **A)** Fitted data after optimized multi-cycle kinetic analysis of corrected responses using a 1:1 binding model. Coloured lines represent original traces for picomolar to nanomolar concentrations of TNFα, with responses that are specific and dose dependent. **B)** Average kinetic parameters calculated from either three or four independent experiments with standard error of mean for TNFα binding to each of three immobilized nanobodies. **C)** Two dimensional iso-affinity kinetic plot of rate constants. Diagonal lines depict equilibrium binding constants and are shown to help with the visualization of the affinity distribution. Each circle represents values determined in an independent run with the mean value labeled in bold.

### In-silico immunogenicity prediction of TNFI-Nb1, TNFI-Nb2, and TNFI-Nb3

To evaluate the immunogenicity risk of TNFI-Nabs, we employed the iTope-AI platform in conjunction with the TCED™ database. iTope-AI is an in-silico immunogenicity risk assessment and deimmunization tool powered by augmented intelligence, utilizing a state-of-the-art machine learning algorithm to predict peptide binding to HLA class II isotypes DR, DP, and DQ. The platform includes 46 common HLA alleles worldwide, ensuring broad population coverage without bias toward any specific ethnic group. Key binding residues were identified through the generation of overlapping 9-mer peptides, each overlapping by eight amino acids, spanning the entire protein sequence. In-house evaluation by Abzena confirmed that iTope-AI accurately predicts 95% of peptide binding core motifs identified by X-ray crystallography and correctly predicts known promiscuous HLA binders, demonstrating high specificity.

iTope-AI assigns binding scores (0-3) to each peptide for each HLA class II allotype, summing these to generate a “Position Risk Score” for each peptide. Peptides are categorized as weak (1-2), medium (3-5), or strong (6+) binders. The “Total Score” for a protein is the sum of all Position Risk Scores, and the highest score (“Hotspot Max”) indicates the presence of strong binders or highly promiscuous peptides. Peptides homologous to human proteome sequences are excluded to account for T cell tolerance. Promiscuous binders are cross-referenced with the TCED™ database to identify potential T cell epitopes.

Although high-affinity MHC class II-binding peptides are associated with T cell epitopes (15, 16), predictive models like iTope-AI do not capture all complexities of antigen presentation, including peptide accessibility, MHC binding stability, and T cell repertoire (17, 18). Consequently, iTope-AI tends to overestimate immunogenic potential by considering all possible peptide iterations, though only a subset is processed and recognized in vivo. This inherent discrepancy highlights the conservative nature of in-silico methods, designed to minimize the risk of overlooking potential immunogenic epitopes.

The iTope-AI analysis identified fourteen non-germline 9-mer MHC class II-binding peptides for TNFI-Nb1, thirteen for TNFI-Nb2, and thirteen for TNFI-Nb3, with corresponding risk scores listed in **Table 3**. Specifically, TNFI-Nb1 achieved a Total Score of 72 and a Hotspot Max of 20, while TNFI-Nb2 and TNFI-Nb3 attained Total Scores of 59 and 62 and Hotspot Max values of 12 and 13, respectively (**Figure 3**). Of the peptides identified for TNFI-Nb1, seven partially matched sequences in the TCED™ database (**Table 4**). Similarly, TNFI-Nb2’s thirteen binders included seven partial matches, and TNFI-Nb3’s binders included eight partial matches to previously identified T cell epitopes (Table 3). The contribution of specific anchor positions to MHC class II binding follows the hierarchy: P1 > P9 > P7 ≥ P6 ≥ P4 (19, 20).

**Table 3.**
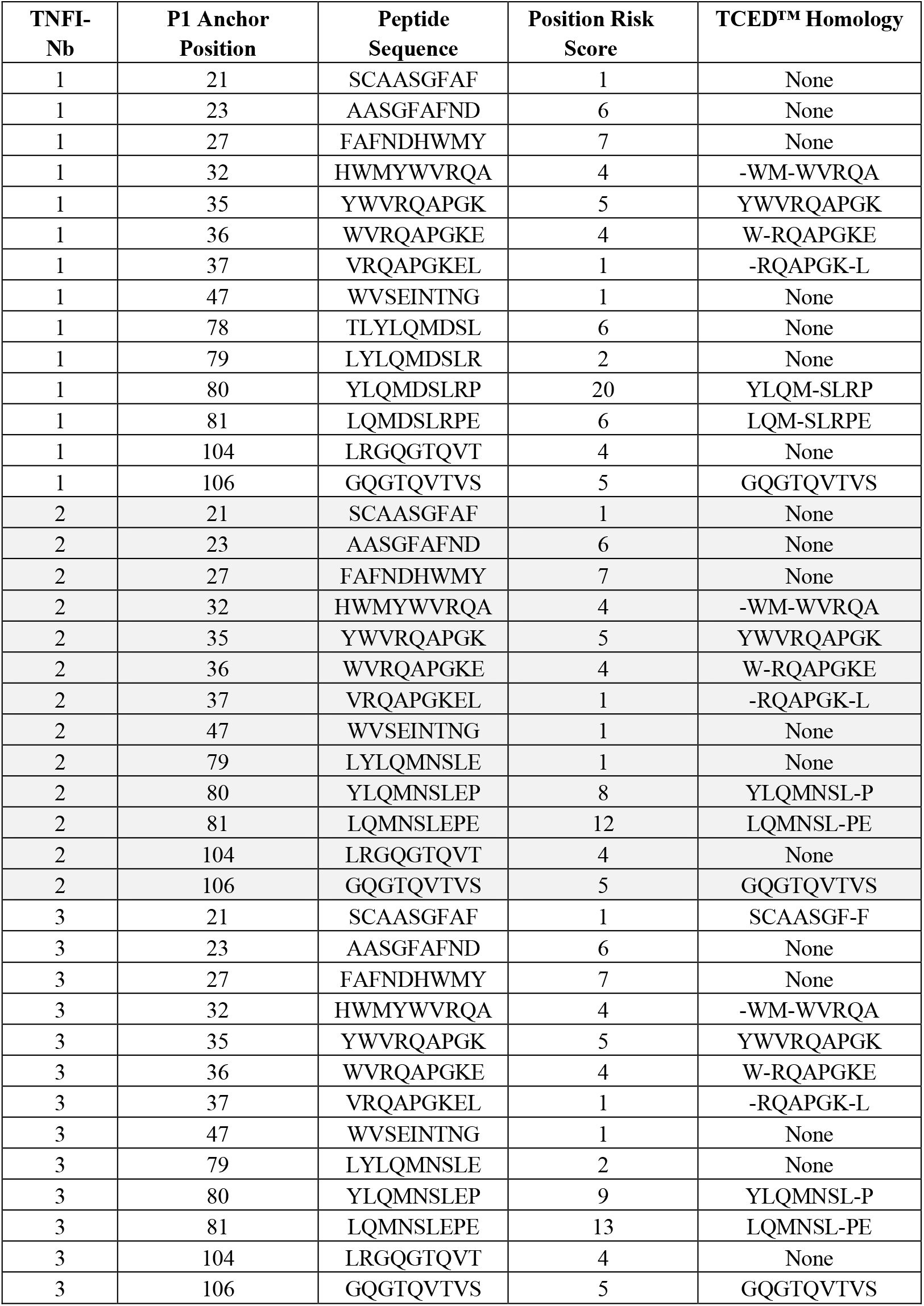
Summary of non-germline 9-mer MHC Class II-binding peptides for TNFI-Nbs exhibiting promiscuous binding. Each peptide is presented with its P_1_ anchor position, amino acid sequence, and Position Risk Score. Homologous peptides identified in the TCED™ database are listed, with mismatched residues indicated by (−).

**Table 4.**
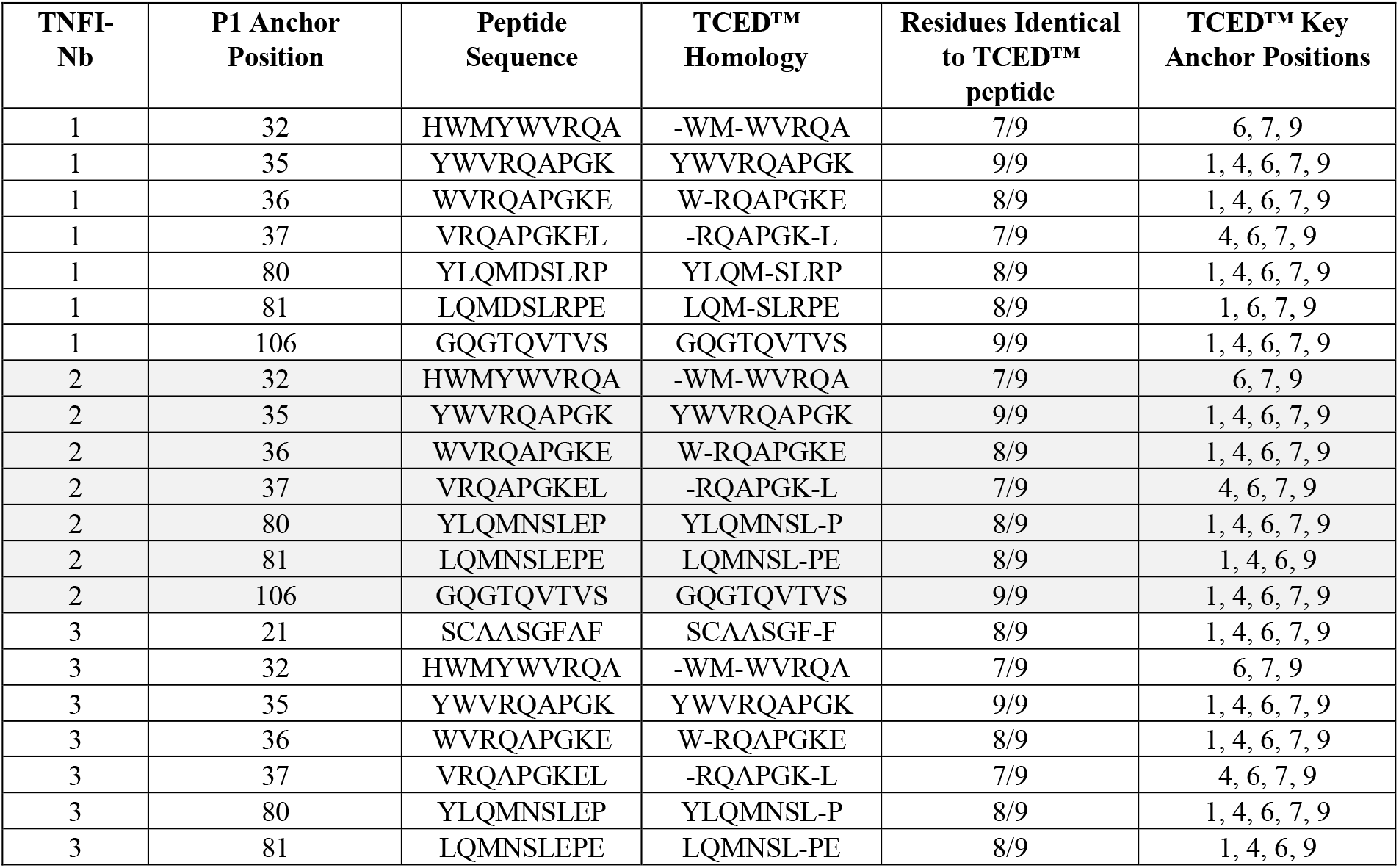
Summary of TCED™ interrogation for non-germline inding peptides of TNFI-Nbs. Homologous peptides identified from the TCED™ database that match the anchor position hierarchy (P_1_ > P_9_ > P_7_ ≥ P_6_ ≥ P_4_) are listed.

**Figure 3.**
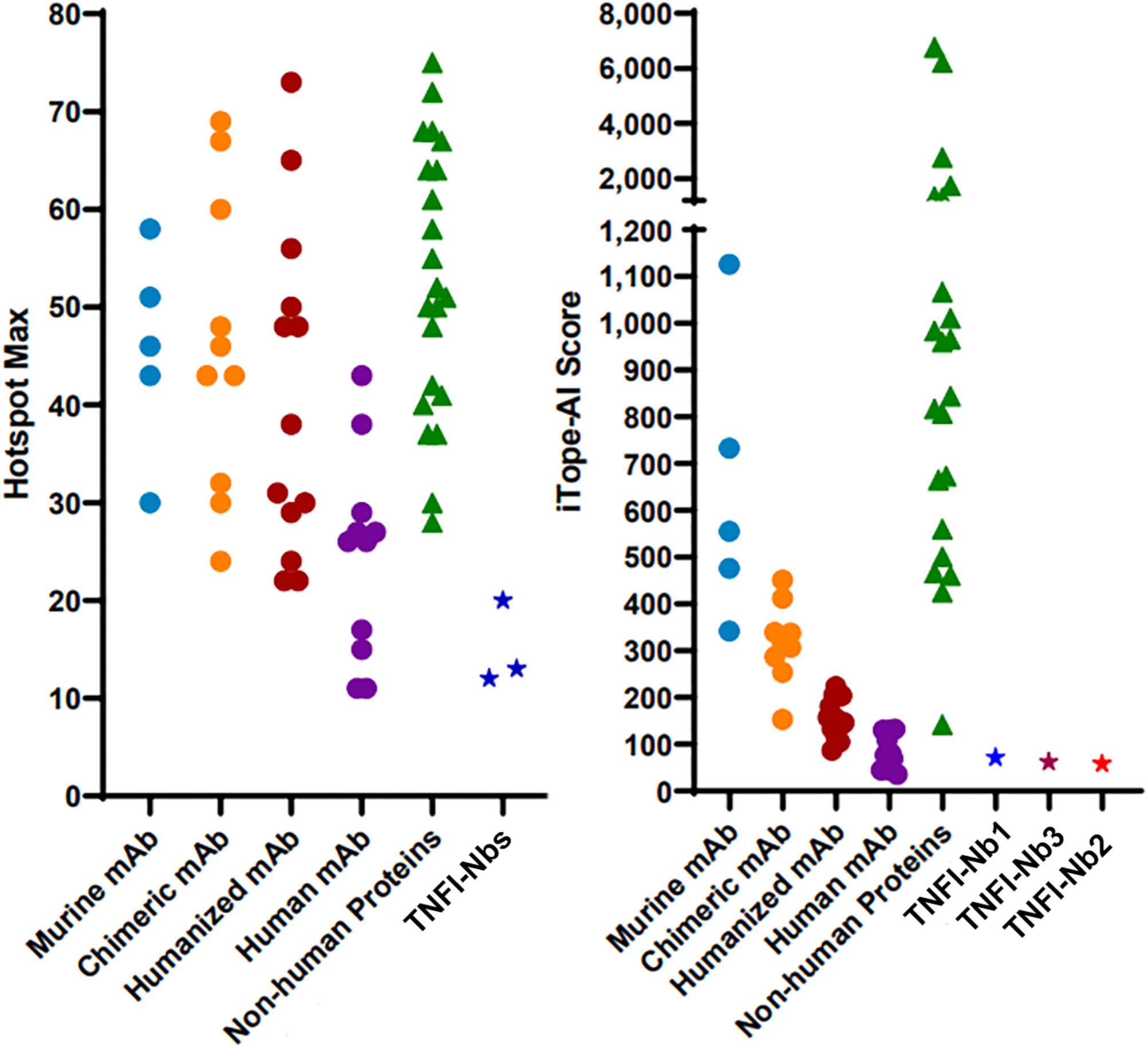
TNFI-Nb1, TNFI-Nb2 and TNFI-Nb3 have low predicted immunogenicity. Total Scores and Hotspot Max for a range of therapeutic antibodies and non-human proteins compared to TNFI-Nb1, TNFINb2 and TNFI-Nb3 indicates low immunogenicity risk for TNFI-Nbs.

A diverse array of therapeutic antibodies and non-human proteins were evaluated by Abzena using the iTope-AI platform, with results illustrated in **Figure 3**. TNFI-Nbs exhibited Total Scores and Hotspot Max values as low as those of the least immunogenic human monoclonal antibodies currently utilized in clinical therapy. In contrast, many of the human monoclonal antibodies assessed displayed scores more than double those of TNFI-Nbs. This comparison highlights the favorable immunogenicity profile of TNFI-Nbs, underscoring their strong potential for therapeutic applications in TNFα-mediated diseases.

### In silico humanization and developability optimization of lead TNFI nanobodies

The nanobody panel, comprising TNFI-Nb1, TNFI-Nb2, and TNFI-Nb3, underwent comprehensive humanization and developability optimization using the machine learning algorithm AbNatiV (21). AbNatiV is a recently introduced AI framework that can accurately predict both humanness and VHH-nativeness of nanobodies from the sequence alone. Humanness scores greater than 0.8 correspond to typically human variable-domain sequences, thereby minimizing potential immunogenicity, while scores below 0.8 indicate non-human origins, which is associated with a higher likelihood of immunogenicity. Among the three nanobodies, TNFI-Nb1 was selected as the optimal starting WT for humanization based on its superior initial humanness and CamSol solubility scores (22), coupled with robust VHH-nativeness (**Table 5**).

**Table 5.** AbNatiV Assessment of TNFI-Nbs. Amino acid residues that differ among the sequences are in bold. Complementarity-determining regions (CDRs), defined by the AHo numbering scheme, are highlighted with gray shading. Key evaluation metrics include Humanness (H), VHH-nativeness (V), and CamSol intrinsic solubility scores (S). Based on the highest humanness and solubility scores, TNFI-Nb1 was selected for the derivation of optimized mutant nanobodies.

| ID | Sequence |  |  |  |
| --- | --- | --- | --- | --- |
| 1 | **EVQLVESGGGLVQPGGSLRLSCAASGFAFNDHWMYWVRQAPGKEL****EWVSEINTNGLITKYADFVKGRFTVSRD****NAKNTLYLQMDSLRPEDTAVYSCSRNQIGTLRGQGTQVTVSS** |  |  |  |
| 2 | **QVQLVESGGGLVQPGGSLRLSCAASGFAFNDHWMYWVRQAPGKEL****EWVSEINTNGLITKYADFVKGRFTVSRD****NAKNTLYLQMNSLEPEDTAVYSCSRNQIGTLRGQGTQVTVSS** |  |  |  |
| 3 | **QVQLVESGGGLVQPGGSLRLSCAASGFAFNDHWMYWVRQAPGKEL****EWVSEINTNGLITKYADFVKGRFTVSRD****NAENTLYLQMNSLEPEDTAVYSCSRNQIGTLRGQGTQVTVSS** |  |  |  |
|  | Name | Humanness | VHH-nativeness | Intrinsic Solubility |
| 1 | 05C08 (TNFI-Nb1) | 0.780 | 0.822 | 0.947 |
| 2 | 05B05 (TNFI-Nb2) | 0.761 | 0.837 | 0.795 |
| 3 | 03H04 (TNFI-Nb3) | 0.755 | 0.838 | 0.807 |

Two distinct mutational sampling strategies were employed: enhanced sampling, which iteratively explores the mutational landscape to rapidly converge on a single humanized sequence, and exhaustive sampling, which systematically assesses all permissible mutation combinations within the position-specific scoring matrices (PSSMs) of human VH and nanobodies (21). Both approaches identify optimal variants on the Pareto Front, maximizing humanness without compromising VHH-nativeness, which is important to retain stability and folding in the absence of a VL domain (21). Importantly, humanization focused exclusively on solvent-exposed residues within the framework regions, and CDR regions were excluded to minimize the risk of hindering binding. We noted that both automated humanization pipelines suggested a mutation in the C-terminal stem of the CDR3 loop, which is an arginine in all TNFI-Nbs (**Table 3**) but is much more commonly a tryptophan in native nanobodies and human VH domains. Consequently, we hypothesized that a rare residue at this stem position next to the CDR3 loop may serve an important functional role, and we thus reverted it to WT arginine in some designs prior to experimental testing (**Table 6**).

**Table 6.**
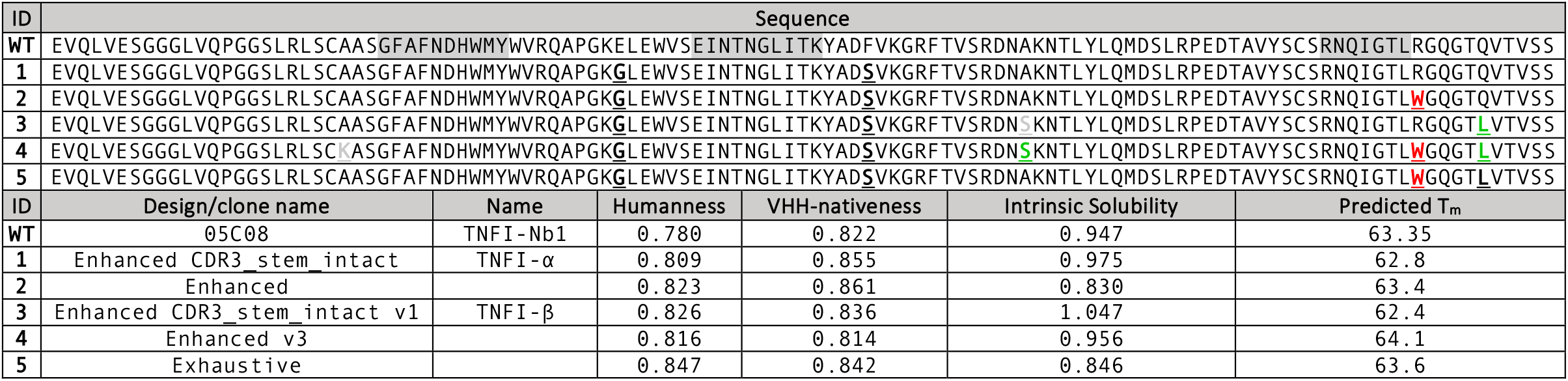
TNFI-Nb Mutants with Enhanced Therapeutic Potential Determined by in-silico Analysis. Mutations are underlined and in bold. Nuerical values in the columns are the Humanness and VHH-nativeness scores predicted by AbNativ, the intrinsic sequence-based solubility score predicted by CamSol, and the apparent melting temperature predicted with NanoMelt. The mutation in red at the stem of the CDR3 region is a mutation that was suggested by both Enhanced and Exhaustive AbNatiV automated sequence-based humanization pipelines. This fact is not surprising as the WT arginine residue is very rare at this position in natural nanobodies, and in fact the mutation to the much more common tryptophan increases both humanness and VHH-nativeness (and predicted melting temperature slightly). However, we hypothesized that a rare residue at the stem of the CDR3 may play an important functional role in binding to the target, and therefore we reverted it to WT in all ‘stem_intact’ designs. Indeed, R is also present at this position in TNFI-Nb2 and 3, further supporting its functional relevance. The mutations in green in designs v1 and v3 were suggested by the CamSol Combination pipeline as mutations predicted to increase solubility and/or stability, without reducing humanness (as assessed by AbNativ).

As a further control for structural integrity post-humanization, the structures of wild-type (WT) and all humanized sequences were modelled with ImmuneBuilder (23). The modelled structures were superimposed based on their framework regions, and the root-mean-square deviation (RMSD) calculated for the CDR regions was approximately 1 Å. This value is significantly smaller than the expected modelling accuracy for these regions(23), suggesting minimal or no displacement of the CDR loops as a result of the humanizing framework mutations.

Following in silico humanization, we used the structural models of both Enhanced and Exhaustive humanized variants as inputs for the CamSol Combination pipeline (24), by excluding all CDR regions from the design and by using an alignment of human VH sequences as input. CamSol Combination automatically identifies combinations of mutations predicted to improve solubility and conformational stability, or one of these properties without affecting the other. The apparent melting temperature of in silico mutants was predicted with NanoMelt (25). Of the mutations suggested by this approach, we retained only those that didn’t reduce humanness according to AbNatiV scoring (**Table 6**).

In summary, 5 TNFI-Nb1 computationally designed variants were selected for experimental characterization (**Table** 6). Three of these had the R to W mutation in the stem of the CDR3 loop, while in the other three this mutation was reverted to reduce the risk of disrupting binding. These designed variants were produced in CHO-S cells through outsourcing to GenScript.

### Assessment of TNFα inhibitory activity in optimized TNFI-Nb1 mutants

To evaluate the TNFI activity of the five TNFI-Nb1 mutants and compare it to that of TNFI-Nb1 on TNFα-induced apoptosis, we employed the IncuCyte Caspase-3/7 Activation Assay using the IncuCyte live-cell imaging system (Sartorius). This assay enables real-time kinetic monitoring of apoptotic events in living cells. WEHI-13VAR cells were treated as described for the cell death assays, with the addition of Caspase-3/7 Green Reagent to each well at a final concentration of 5 μM. This fluorogenic substrate specifically detects activated caspase-3 and caspase-7, key executors of apoptosis. Plates were placed in the IncuCyte system, which maintained optimal incubation conditions while capturing fluorescence images at 30-minute intervals. In these experiments, TNFI-Nbs were tested using 2-fold serial dilutions ranging from 10 nM to 12.2 pM. The 10-hour time point was selected to assess TNFI activity because caspase-3/7 activity induced by Actinomycin D alone was minimal, whereas activity induced by TNFα plus Actinomycin D was significantly elevated. This contrast allowed for more accurate measurements of the nanobodies’ effects on TNFα-dependent caspase-3/7 activation. The IncuCyte software quantified fluorescence intensity, correlating to caspase-3/7 activity and thereby apoptotic cell death. Data were normalized to control wells treated with Actinomycin-D alone. The concentration of each nanobody required to achieve 50% inhibition of caspase-3/7 activation (IC_50_), compared to the wells treated with TNFa plus Actinomycin D only, was calculated.

The IncuCyte Caspase-3/7 activation assays revealed varying inhibitory activities among the tested nanobody mutants against TNFα-induced caspase-3/7 activation. The Enhanced, Enhanced v3, and Exhaustive mutants exhibited IC_50_ values of 1.417 nM, 29.896 nM, and 2.27 nM, respectively, indicating reduced potency compared to the parental TNFI-Nb1 (IC_50_ = 133.5 pM). This reduction in potency is likely attributable to the W to R mutation at the stem of the CDR3 region. In contrast, the Enhanced CDR3_stem_intact and CDR3_stem_intact v1 mutants, which lack this mutation, demonstrated significantly improved IC_50_ values of 48.84 pM and 64.31 pM, respectively, reflecting enhanced inhibitory activity against TNFα. For simplicity, Enhanced CDR3_stem_intact and CDR3_stem_intact v1 have been renamed TNFI-α and TNFI-β, respectively (Figure 4A and 4B). These optimized mutants, TNFI-α and TNFI-β, exhibit superior inhibitory performance, underscoring their strong potential for clinical development by offering increased efficacy alongside a reduced likelihood of anti-drug antibody (ADA) formation.

**Figure 4.**
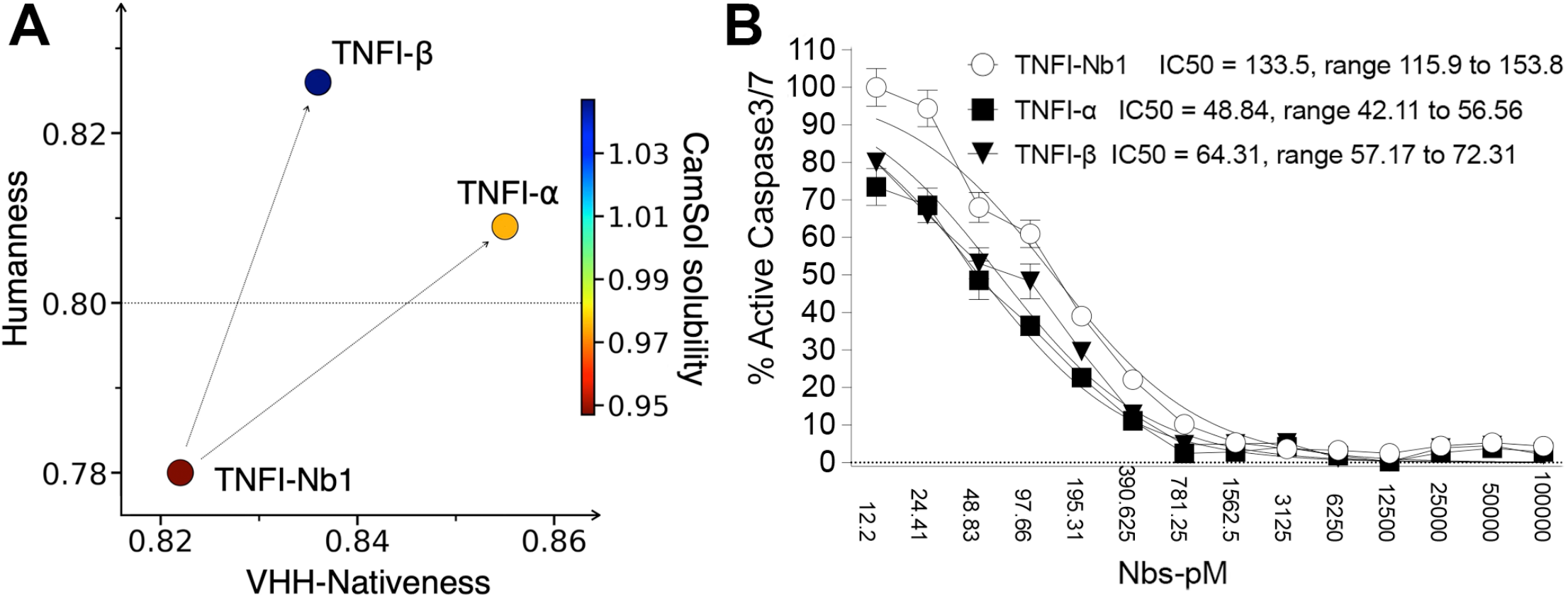
In silico engineering of TNFI-α and TNFI-β nanobodies: optimized mutants of TNFI-Nb1 with enhanced TNFI activity. **A**) Scatter plot with the AbNatiV VH humanness score (y-axis) as a function of the AbNatiV VHH-Nativeness score (x-axis), showing the predicted improvements of TNFI-α and TNFI-β over the WT TNFI-Nb1. Points are colored according to their CamSol Intrinsic solubility score (color bar). The horizontal line at 0.8 humanness represents the threshold that best separates human (humanness > 0.8) from non-human (humanness < 0.8) VH sequences. **B**) WEHI-13VAR cells were treated for 10 hours with 0.25 ng/mL active trimeric human sTNFα (supplemented with 1 μg/mL Actinomycin-D, Sigma), either alone or in combination with 2-fold serial dilutions of TNFI-Nbs, ranging from 10,000 pM to 12.2 pM. Apoptosis was measured using the iQue® Caspase 3/7 Reagent Kit (Sartorius), which employs a fluorogenic dye that fluoresces upon cleavage by activated caspases 3 and 7, essential mediators of the apoptotic pathway. The IC50 values represent the concentration of each TNFI-Nb required to inhibit 50% of TNFα-induced Caspase 3/7 activation. Data illustrate the enhanced inhibitory activity of TNFI-α and TNFI-β mutants compared to TNFI-Nb1.

### SPR analysis of TNFI-α and TNFI-β binding to human TNFα

To evaluate the binding affinities of TNFI-α and TNFI-β to human TNFα, SPR analysis was performed. The purity of the nanobody samples, TNFI-α and TNFI-β, along with human TNFα, was confirmed to exceed 90%, ensuring high-quality reagents for subsequent analyses. Assay development and kinetic analysis were conducted using Cytiva’s Biacore 1K SPR system with Carboxyl CM5 sensors. This process involved optimizing immobilization conditions, buffer formulations, regeneration protocols, and kinetic fitting models to establish reliable and reproducible measurement conditions. Optimal immobilization was achieved at a ligand concentration of 2.5 μg/mL in 10 mM acetate buffer (pH 4.5), resulting in response maxima (Rmax) of 34.44 RU for TNFI-α and 12.74 RU for TNFI-β, aligning with theoretical predictions. Buffer optimization identified 1x PBS supplemented with 3 mM EDTA and 0.05% v/v Surfactant P20 as the optimal running buffer, effectively minimizing non-specific binding and maximizing signal-to-noise ratios.

Kinetic analysis revealed that the interactions between TNFα and both nanobodies adhered to a 1:1 binding model (Figure 5A), with equilibrium dissociation constants (KD) of 167 pM for TNFI-α and 405 pM for TNFI-β (Figure 5B), indicating high affinity binding. Iso-affinity analysis further demonstrated that TNFI-α exhibited a significantly faster association rate (ka) compared to TNFI-β, contributing to its superior overall binding affinity (Figure 5C). These results indicate that TNFI-α and TNFI-β exhibit high affinity for TNFα, comparable to the affinities observed for TNFI-Nb1, TNFI-Nb2, and TNFI-Nb3, underscoring the potential of TNFI-α and TNFI-β as highly effective therapeutic agents targeting TNFα-mediated pathways. The SPR experiments were outsourced to Rapid Novor.

**Figure 5.**
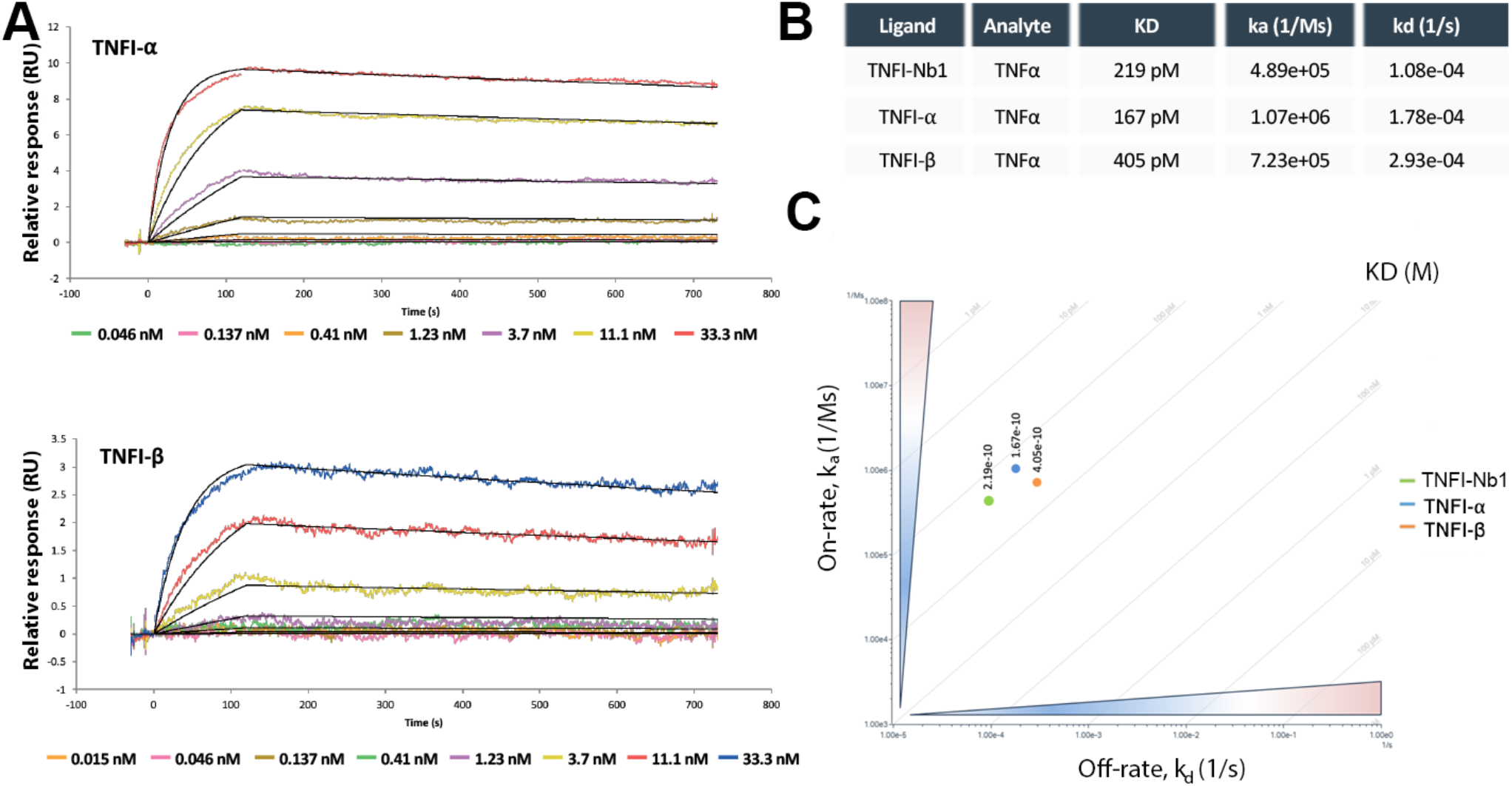
Kinetics of TNFα analytes binding to immobilized nanobodies. Kinetic analysis of TNFα binding to humanized nanobodies TNFI-α and TNFI-β directly immobilized on CM5 Carboxyl sensors via amine coupling. **(A)** Fitted data after optimized multi-cycle kinetic analysis of corrected responses using a 1:1 binding model. Coloured lines represent original traces for nanomolar concentrations of TNFα, with responses that are specific and dose dependent. **B)** Kinetic parameters were determined, characterizing kinetics of TNFα binding to immobilized TNFI-α and TNFI-β nanobodies. **C)** Two dimensional iso-affinity kinetic plot of rate constants. Diagonal lines depict equilibrium binding constants and are shown to help with the visualization of the affinity distribution. Each circle represents one set of kinetic values determined based on a 1:1 binding model.

As summarized in Table 5, the humanization process yielded two nanobodies with enhanced humanness scores without compromising their biological activity, specifically TNFα inhibition. In fact, the activity of the humanized nanobodies appears to be slightly enhanced. TNFI-α demonstrated the lowest IC50 value (48.84 pM) compared to TNFI-β (64.31 pM), indicating superior inhibitory activity. However, TNFI-β exhibited higher humanness (0.826 vs. 0.809) and solubility (1.047 vs. 0.975) scores. These complementary strengths make both TNFI-α and TNFI-β strong candidates for further development.

TNFI-α and TNFI-β, respectively have been assigned by NanoNewron commercial designations as follows: TNFI-α: NN-223; TNFI-β: NN-224. Both NN-223 and NN-224 are currently under evaluation for further preclinical development. These variants represent promising candidates for therapeutic applications targeting human TNFα.

## Discussion

In this study we present the generation, characterization, and optimization of TNFI-Nbs NN-223 and NN-224. Through humanization and functional optimization, NN-223 (formerly TNFI-α) and NN-224 (formerly TNFI-β) emerged as lead candidates, exhibiting enhanced predicted humanness and solubility coupled with slightly improved measured TNFα inhibitory activity. These findings underscore the efficacy of nanobody engineering in producing biologics with superior therapeutic profiles. For clarity, these nanobodies will be referred to by their commercial names, NN-223 and NN-224, throughout the remainder of this discussion.

The manufacturing process for these nanobodies, achieved through transient transfection in CHO-S cells, yielded high purity and very low endotoxin levels. These attributes indicate that the production process is scalable for clinical applications.

SPR analysis confirmed that both NN-223 and NN-224 maintain high binding affinities for soluble human TNFα, with KD values of 167 pM and 405 pM, respectively. These affinities are comparable to those of earlier TNFI-Nb prototypes (TNFI-Nb1, TNFI-Nb2, and TNFI-Nb3). The ability of NN-223 and NN-224 to retain high affinity following humanization demonstrates the effectiveness of framework optimization strategies in minimizing immunogenic potential while preserving critical antigen-binding regions.

Functional assays assessing TNFα-induced caspase-3/7 activation revealed significantly improved IC_50_ values for NN-223 and NN-224 compared to TNFI-Nb1. Specifically, NN-223 exhibited the most potent inhibitory activity with an IC_50_ of 48.84 pM, while NN-224 achieved an IC_50_ of 64.31 pM. These enhanced inhibitory efficacies, combined with favorable production parameters, bolster the clinical development potential of both nanobodies by offering superior efficacy alongside a reduced likelihood of anti-drug antibody (ADA) formation.

The therapeutic efficacy of TNFIs in diseases like rheumatoid arthritis, Crohn’s disease, and psoriasis illustrates the ability of TNFα inhibition to significantly reduce systemic inflammation, even in conditions where other cytokines contribute to the inflammatory milieu (1-5). This highlights TNFα’s central role in modulating global inflammatory responses. Unfortunately, FDA-approved TNFIs biologic, the most potent and specific TNFIs, have limited BBB permeability (CSF/Serum ratios ∼0.001) restricting their use in therapy of neuroinflammatory disorders, such as AD (26-29).

Emerging evidence from both human and animal studies implicates TNFα as a pivotal factor in AD pathogenesis (30, 31). Several findings highlight TNFα’s role and its potential as a therapeutic target in AD.

### Human Evidence

Elevated levels of TNFα have been consistently observed in the blood, cerebrospinal fluid (CSF), and CNS of AD patients (32-36). These elevated TNFα levels are associated with increased neuroinflammation and are implicated in disease progression through both inflammatory and non-inflammatory mechanisms.

Genetic studies have identified polymorphisms in TNFα and its receptors that are associated with increased AD risk (37, 38), suggesting that these variants may modulate TNFα expression or receptor activity, thereby amplifying its pathological impact.

Microglia, the primary producers of TNFα in the CNS, play a crucial role in AD pathogenesis (33, 39). Genetic studies have linked microglia-specific genes, such as TREM2, to late-onset AD (LOAD) (40, 41). Pathogenic variants like TREM2 p.R47H impair essential microglial functions, including phagocytosis and regulation of TNFα production, thereby exacerbating neuroinflammation and neuronal damage.

Clinical evidence supports the therapeutic potential of TNFα inhibition. Patients with inflammatory diseases treated with TNFIs exhibit a reduced risk of developing AD (42, 43). Additionally, peri-spinal and intrathecal administration of TNFIs, such as etanercept and infliximab, has shown cognitive improvements in sporadic AD cases (44, 45). These observations suggest that TNFα inhibition can mitigate neuroinflammation and associated cognitive decline in AD.

### Mechanistic insights from model organisms and in vitro studies

Elevated TNFα levels promote amyloidbeta (Aβ) production and impair its clearance, primarily through microglial dysfunction (46). Deletion of TNF receptor 1 (TNFR1) in AD mouse models reduces Aβ formation and alleviates learning and memory deficits (47). Furthermore, TNFα regulates synaptic receptor trafficking by promoting AMPA receptor export and GABA receptor import. Supraphysiological TNFα disrupts this balance, favoring excitatory dominance, impairing long-term potentiation (LTP), and increasing excitotoxicity risk (48-50). As Aβ production and tau secretion are activity-dependent processes (51-53), TNFα-driven excitatory activity may exacerbate both Aβ and tau pathology.

Data from *Trem2*^*R47H*^ knock-in (KI) rats, a model for Late Onset AD, provide support for TNFα’s role in AD pathogenesis. These KI rats exhibit early and significant increases in TNFα levels in the CNS and CSF. Elevated TNFα disrupts the excitatory/inhibitory balance, increases excitatory transmission, and reduces inhibitory transmission, thereby impairing LTP and exacerbating excitotoxicity. Administration of low doses of an anti-TNFα antibody with TNFI activity reverses these deficits, restoring excitatory/inhibitory balance, normalizing LTP, and mitigating the harmful effects of TNFα dysregulation (54-56).

### Advantages of nanobodies over traditional biologic TNFIs

Unlike larger TNFI biologics, the smaller size and robust structural stability of nanobodies may facilitate better BBB penetration (8, 9), enhancing their therapeutic reach within the brain. Furthermore, nanobodies can be engineered for improved BBB permeability through strategies such as receptor-mediated transcytosis via Transferrin receptor 1 (10-14), potentially increasing their efficacy in treating neurodegenerative disorders. Thus, future studies will focus on evaluating the in vivo efficacy of NN-223 and NN-224, their pharmacokinetics, and their ability to cross the BBB, which would enable their use in CNS diseases linked to TNFα dysregulation. Comparative analyses in preclinical models will also help determine their relative suitability for clinical development based on efficacy, pharmacokinetics, and immunogenicity.

*In conclusion*, NN-223 and NN-224 represent significant advancements in nanobody engineering, offering high specificity, strong inhibitory activity, and favorable developability profiles. These optimized nanobodies demonstrate their potential as next-generation therapeutics for diseases driven by dysregulated TNFα signaling, including CNS inflammatory disorders such as AD. The combination of enhanced inhibitory efficacy, scalable manufacturing processes, and reduced immunogenic potential positions NN-223 and NN-224 as promising candidates for clinical development.

## Experimental procedures

### Antibodies and reagents

WEHI-13VAR cells (ATCC, CRL-2168) and HEK293T cells (ATCC, CRL-3216) were utilized in this study. Detailed information for the mammalian expression plasmids for Human TNFα and EGFP (VB220203-1058zds) can be found here https://en.vectorbuilder.com/vector/VB220203-1058zds.html. The anti-His tag APC-conjugated antibody was purchased from R&D Systems (Catalog No. IC050A). Fugene transfection reagent was acquired from Promega (Catalog No. E2311). Recombinant human TNFα protein was sourced from Acro Biosystems (Catalog No. TNA-H5228), recombinant mouse TNFα protein from Acro Biosystems (Catalog No. TNA-M82E9), and recombinant rat TNFα protein from R&D Systems (Catalog No. 510-RT-010/CF). Fetal bovine serum was obtained from Gibco (Catalog No. A3840102). EDTA was purchased from Sigma-Aldrich (Catalog No. N6507). Propidium iodide was acquired from Invitrogen (Catalog No. P3566). DMEM and RPMI 1640 media were purchased from Corning (Catalog Nos. 10-017-CV and 10-040-CV, respectively). Acridine orange/propidium iodide (AO/PI) staining solution was obtained from DeNovix (Catalog No. CD-AO-PI-7.5). Cell Counting Kit 8 (CCK-8) was sourced from DojinDo (Catalog No. CK-04). Actinomycin-D was acquired from Sigma-Aldrich (Catalog No. A9415), and Caspase-3/7 green dye was purchased from Thermo Fisher Scientific (Catalog No. C10423).

### Generation and screening of camelid anti-TNFα nanobodies

To generate anti-TNFα nanobodies (a-TNFα-Nabs), one alpaca and one llama were immunized with active trimeric human TNFα (Acro Biosystems, TNA-H5228). The immunization protocol began with an initial subcutaneous injection of 0.5 mg TNFα mixed with Complete Freund’s Adjuvant (CFA) at week 0, followed by booster injections of 0.5 mg TNFα with Incomplete Freund’s Adjuvant (IFA) every two weeks up to week 14. Serum samples were collected before immunization and at designated time points to assess antibody titers via ELISA, utilizing TNF-alpha-coated plates with appropriate negative and positive controls to ensure specificity.

At weeks 10 and 14, 500 mL of whole blood were drawn from each llama for peripheral blood mononuclear cell (PBMC) isolation. PBMCs were isolated within four hours of blood collection using density gradient centrifugation, ensuring cell viability above 99% as determined by trypan blue exclusion. Total RNA was extracted from the isolated PBMCs using the RNeasy Maxi Kit (Qiagen) and quantified by spectrophotometry, ensuring an A260/A280 ratio greater than 1.9. High-quality RNA was confirmed by agarose gel electrophoresis, displaying distinct 18S and 28S rRNA bands without signs of degradation.

Complementary DNA (cDNA) was synthesized from the purified RNA using the SuperScript IV FirstStrand Synthesis System (Thermo Fisher Scientific).

A nanobody-specific library was constructed by amplifying the variable regions of heavy-chain-only antibodies (VHH) through a two-step PCR process using camelid-specific degenerate primers. The amplified VHH fragments were cloned into the pADL-20c phagemid vector using SfiI restriction sites and transformed into E. coli TG1 cells, resulting in a phage display library containing approximately 2.57 × 10^9^ individual clones. Library diversity was confirmed by sequencing 74 random clones, which revealed 88% contained VHH inserts with intact open reading frames and no duplicate sequences.

Library panning was performed against immobilized TNFα through three rounds of selection to enrich for specific binders. To reduce non-specific interactions, the phage pool was pre-absorbed on BSA-coated wells before each panning round. Enrichment of specific phage binders was monitored using dot assays, which demonstrated increased binding to TNFα with each successive round.

Following panning, 94 individual clones were screened using an off-phage ELISA to identify those with specific binding to TNFα and minimal binding to BSA. Positive clones were further validated through repeat screening and sequencing to ensure specificity and diversity. Selected nanobodies were expressed in a non-amber-suppressor strain of E. coli, and periplasmic fractions containing His-tagged nanobodies were purified using His-tag affinity chromatography. Purity was confirmed by SDS-PAGE, and nanobody concentrations were determined by absorbance at 280 nm. Purified nanobodies were dialyzed into PBS (pH 7.4) and filter-sterilized for downstream applications.

### Production of nanobodies in CHO-S cells

Protein production was outsourced to GenScript, and detailed experimental procedures and results are provided in the Supporting information data files entitled: TNFI-Nb Production in CHO-S cells and Optimized TNFI-Nbs Production.

### SPR analyses

These analyses were outsourced to Rapid Novor. The purity of the nanobodies as well as human TNFα, was confirmed to exceed 90%, ensuring high-quality reagents for subsequent assays. Kinetic analyses were performed using Cytiva’s Biacore 1K SPR system and Nicoya’s OpenSPR-XT instrument with Carboxyl CM5 sensors. Details of experimental procedures are provided in the Supporting information data files entitled: SPR Analysis of TNFI-Nb1-3 Binding to Human TNFα and SPR Analysis of Optimized Mutants TNFI-α and TNFI-β Binding to Human TNFα.

### In-silico immunogenicity prediction

Immunogenicity risk of TNFI-Nabs was evaluated employing the iTope-AI platform in conjunction with the TCED™ database. The analysis was performed by Abzena. Details of experimental procedures are provided in the Supporting information data files entitled: iTope analysis of TNFI-Nb1-3 immunogenicity.

### TNFI Activity and IC_50_ Determination Assays

WEHI 13VAR cells were cultured in RPMI 1640 medium supplemented with 10% fetal bovine serum (FBS) in a humidified incubator maintained at 37°C with 5% CO_2_. Cells were harvested, and cell number and viability were assessed using acridine orange/propidium iodide (AO/PI) staining with a Denovix Celldrop cell counter. The cell suspension was adjusted to a density of 30,000 cells per well, and 90 μL of this suspension was seeded into a 96-well plate containing RPMI 1640 medium with 10% FBS and 1 μg/mL Actinomycin-D. Nanobodies were pre-mixed at serial concentrations with human TNF-α active trimer, mouse TNF-α active trimer, or rat TNF-α active trimer and incubated at room temperature for 30 minutes. Subsequently, 10 μL of the pre-mixed solution was added to each well containing WEHI 13VAR cells, resulting in a final TNF-α concentration of 0.25 ng/mL and the desired final concentrations of Nbs.

The cytotoxicity assay was conducted by incubating the plates for 24 hours at 37°C with 5% CO_2_. Following incubation, 10 μL of Cell Counting Kit 8 (CCK-8) was added to each well, and the plates were incubated for an additional 2–4 hours. Absorbance was measured at 450 nm using a microplate reader. To quantify TNFα-induced cytotoxicity, cytotoxicity values from wells treated with Actinomycin-D alone were subtracted from those treated with both Actinomycin-D and TNFα. The percentage protection conferred by TNFI-Nbs was calculated by setting the cytotoxicity induced by Actinomycin-D plus TNFα as 100%, and expressing the cytotoxicity of samples treated with Actinomycin-D, TNFα, and TNFI-Nbs as a percentage of this reference value.

For the Caspase-3/7 activity assays, cells were treated as described above with one modification: Caspase-3/7 dye was added to the pre-mixed solution containing TNFα, and nanobodies to achieve a final concentration of 0.5 μM upon addition to the cells. The plate was monitored using the IncuCyte live-cell analysis system, maintaining incubation at 37°C with 5% CO_2_. To quantify TNFα-induced Caspase-3/7 activity, activity values from wells treated with Actinomycin-D alone were subtracted from those treated with both Actinomycin-D and TNFα. The percentage protection conferred by TNFI-Nbs was calculated by setting the Caspase-3/7 activity induced by Actinomycin-D plus TNFα as 100%, and expressing the Caspase-3/7 activity of samples treated with Actinomycin-D, TNFα, and TNFI-Nbs as a percentage of this reference value.

### FACS analysis

HEK293T cells were cultured in DMEM medium supplemented with 10% fetal bovine serum (FBS) in a humidified incubator maintained at 37°C with 5% CO_2_. HEK293T cells were transiently transfected with constructs encoding human TNFα and EGFP using Fugene transfection reagent and incubated for 24 hours. Transfected cells were harvested and resuspended in ice-cold FACS buffer (PBS supplemented with 1% BSA and 1 mM EDTA, pH 7.2). Nanobodies were added to the cells in a 96-well plate and incubated at 4°C with shaking for 45 minutes. Following incubation, cells were centrifuged at 1000×g for 5 minutes and washed three times with 200 μL of FACS buffer. The cell pellet was then resuspended in 100 μL of FACS buffer containing anti-His tag APC-conjugated antibody at a 1:100 dilution and incubated for 30 minutes at 4°C with shaking. After another centrifugation at 1000×g for 5 minutes and three additional washes with 200 μL of FACS buffer, the cell pellet was resuspended in 200 μL of FACS buffer containing propidium iodide (PI) at a 1:1000 dilution for subsequent analysis.

### GenBank

The cDNA and amino acid sequences of the nanobodies have been deposited in GenBank under the following accession numbers:

- NN-224/TNFI-β: PV016701
- NN-223/TNFI-α: PV016702
- 05C08R3/TNFI-Nb1: PV016703
- 05B05R3/TNFI-Nb2: PV016704
- 03H04R3/TNFI-Nb3: PV016705

## Statistical Analysis

Data were analyzed using GraphPad Prism 10 software. IC_50_ values were determined using a nonlinear regression model (“log(inhibitor) vs. response”). Differences between groups were assessed using one-way ANOVA followed by Tukey’s post hoc test. A p-value of <0.05 was considered statistically significant.

